# Intracranial-EEG evidence for medial temporal pole driving amygdala activity induced by multi-modal emotional stimuli

**DOI:** 10.1101/2020.02.24.963801

**Authors:** Saurabh Sonkusare, Vinh Thai Nguyen, Rosalyn Moran, Johan van der Meer, Yudan Ren, Nikitas Koussis, Sasha Dionisio, Michael Breakspear, Christine Guo

## Abstract

The temporal pole (TP) is an associative cortical region required for complex cognitive functions such as social and emotional cognition. However, functional mapping of the TP with functional magnetic resonance imaging is technically challenging and thus understanding of its interaction with other key emotional circuitry, such as the amygdala, remain elusive. We exploited the unique advantages of stereo-electroencephalography (SEEG) to assess the responses of the TP and the amygdala during the perception of emotionally salient stimuli of pictures, music and movies. These stimuli consistently elicited high gamma responses (70-140 Hz) in both the TP and the amygdala, accompanied by functional connectivity in the low frequency range (2-12 Hz). Computational analyses suggested the TP driving this effect in the theta-alpha frequency range and which was modulated by the emotional salience of the stimuli. Of note, cross-frequency analysis indicated the phase of theta-alpha oscillations in the TP modulated the amplitude of high gamma activity in the amygdala. These results were reproducible with three types of stimuli including naturalistic stimuli suggesting a hierarchical influence of the TP over the amygdala in non-threatening stimuli.

## Introduction

Human brains are remarkable for their relative size, as well as by a greater expansion of association cortices compared to other mammals (Van Essen and Dierker 2007, Glasser, Sotiropoulos et al. 2013). These association cortices play key roles in conceptual and executive functions. One such association area is the temporal pole (TP, also referred to as BA 38 or area TG) covering the most anterior extent of the temporal lobe (Olson, Plotzker et al. 2007). It is a complex cortical region particularly observed in primate brains, however, its functional contribution to cognition is elusive and lacks consensus.

A prominent theory advances the TP to be a ‘hub’ that integrates conceptual information from different modalities to form domain-general semantic representations (McClelland and Rogers 2003, Rogers, Ralph et al. 2004, Patterson, Nestor et al. 2007, Lambon Ralph and Patterson 2008, Ralph, Jefferies et al. 2017). That is, the TP serves as a transmodal hub, or “conceptual store”, that binds information from different sensory modalities. Functional neuroimaging with PET and MRI supports its transmodal functional role with studies finding TP activation across diverse tasks in different modalities involving auditory, olfactory, memory, vestibular, visual, and linguistic processing (Ardila, Bernal et al. 2014). Other theories have proposed the TP to be a region specialised for social and emotional processing, (Olson, McCoy et al. 2013). However, socio-emotional knowledge can itself be considered a type of semantic memory and conceptual processing (Olson, McCoy et al. 2013).

In contrast to the TP, the amygdala is a subcortical structure present almost ubiquitously across the animal kingdom, even in fish and amphibians (as part of amygdaloid complex) (Moreno and González 2007). It has been extensively studied in many species over recent decades, particularly in mice, where it is crucial in fear conditioning (LeDoux, Cicchetti et al. 1990, Phillips and LeDoux 1992, LeDoux 2003). Amygdala lesion studies in primates (Klüver 1937) and humans (Pilleri 1966, Marlowe, Mancall et al. 1975) also suggest its important role in processing negative emotions. Neuroimaging has provided substantial evidence for the involvement of the amygdala in processing of emotions (positive as well as negative) (LaBar, Gatenby et al. 1998, Morris, Friston et al. 1998, Liberzon, Phan et al. 2003, Critchley 2005, Koelsch, Fritz et al. 2006, Williams, Kemp et al. 2006, Costafreda, Brammer et al. 2008, Motzkin, Philippi et al. 2015, Diano, Tamietto et al. 2017).

Both the TP and the amygdala contribute to emotional processes (Klasen, Weber et al. 2011, Mathiak, Klasen et al. 2011, Pehrs, Zaki et al. 2015, Saarimäki, Gotsopoulos et al. 2015), and are strongly anatomically connected (Bach, Behrens et al. 2011, Gschwind, Pourtois et al. 2011, Abivardi and Bach 2017). However, no study has systematically explored their interactions as a functional dyad. This is partly attributable to technical challenges with conventional brain recording techniques. Specifically, both these regions have been known to be affected by fMRI signal drop-out (Devlin, Russell et al. 2000, Boubela, Kalcher et al. 2015), and low fidelity of EEG signal due to their deep location in the brain (Tesche, Karhu et al. 1996, Riggs, Moses et al. 2009). Here, we sought to comprehensively investigate the interaction between the TP and the amygdala in emotional processing by leveraging the unique advantages of stereo-electroencephalography (sEEG), namely its excellent spatio-temporal resolution. To induce affective states, we employed distinct types of emotional stimuli including conventional static images and naturalistic stimuli (music and movie clips). Naturalistic stimuli, not only offer better ecological validity (Hasson 2004, Sonkusare, Breakspear et al. 2019) but also evoke robust physiological changes thought to be integral to emotional experiences (Gross and Levenson 1995, Seth and Friston 2016). Furthermore, we sought to determine whether responses and interaction between the TP and the amygdala to simplified stimuli generalize to naturalistic conditions.

We anticipated stimulus induced responses in the TP and amygdala, and a functional coupling between them. Due to its role as a conceptual hub and thus in providing contextual information (Pehrs, Zaki et al. 2015), we anticipated that the TP may have a superordinate influence over amygdala activity. We investigated this hypothesis by employing directed functional and effectivity connectivity measures.

## Materials and Methods

### Participants

Participants were drawn from a cohort of 13 patients with medically intractable epilepsy, implanted with stereotactic electrodes for clinical evaluation at Mater advanced Epilepsy Unit, Mater Hospital Brisbane. The choice of regions for electrode implantation was based on clinical criteria alone. Inclusion criteria for this study were the location of electrodes within the temporal pole and the amygdala. 6 patients (1 female; age 17 - 56 years) met these criteria and hence participated in this study. All participants (P2, P4, P8, P10, P13, and P15) had medically intractable epilepsy. Patient characteristics are provided in *Table 1*. Participants gave written informed consent to participate in the study and were free to withdraw from the study at any time. The study was approved by the Human Research Ethics Committees of the Mater Hospitals and the QIMR Berghofer Medical Research Institute and performed in agreement with the Declaration of Helsinki.

**Table 1.**
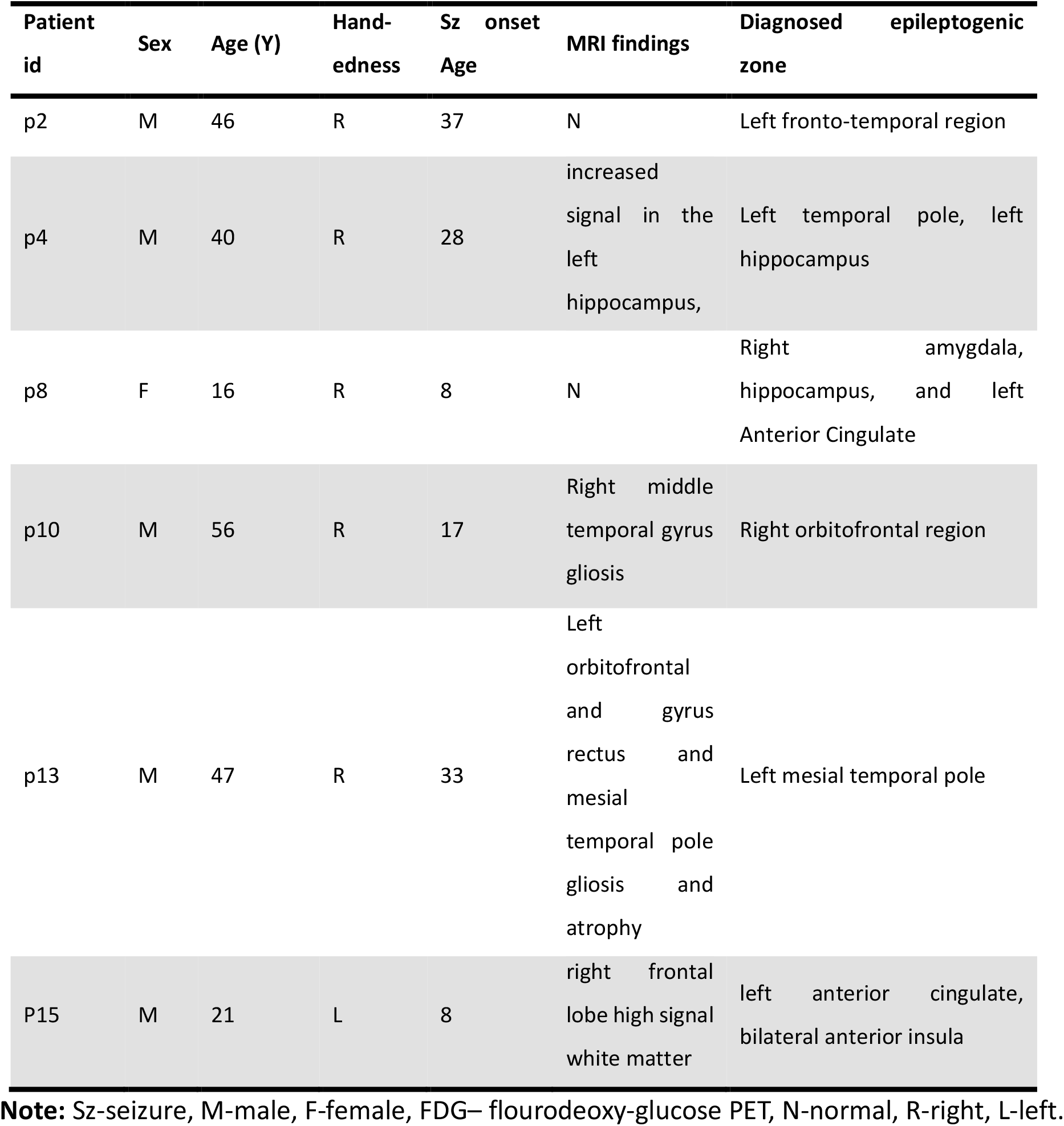
Patient profile.

### Data acquisition

#### Stereo-EEG recordings

Implantation of intracerebral electrodes with multiple contacts (DIXI, 10-15 contacts, electrode diameter: 1 mm, intercontact spacing 1.5 mm) was performed with a stereotactic procedure and planned individually based on the likely seizure onset zone inferred by the epilepsy clinician (author SD) from non-invasive pre-operative studies. sEEG signals sampled at 1 kHz were recorded on a Neurofax EEG-1200 system (Nihon Koden, Japan). All experimental task data were processed off-line using EEGLAB (Delorme and Makeig 2004) and custom routines programed in MATLAB (MathWorks, MA, USA). Recordings were down-sampled to 500 Hz and band-passed filtered between 0.5 to 195 Hz using a zero-phase lag filter (FIR). Power line 50 Hz noise and its harmonics were notch filtered. Data were referenced to common average reference computed from the average of all the clean intra-cranial electrode channels.

### Data analysis

For picture stimuli data, a 500 ms window of data following each stimulus was used. All epochs were visually inspected and those with inter-ictal spikes, defined as paroxysmal discharges lasting less than 250 ms, (Staley, White et al. 2011) were excluded from further analysis. For music clips data, to avoid edge effects at the termination of each clip, only the first 12 s were employed. For movie clips, data corresponding to annotations of most emotional continuous 10 seconds, (as rated by human subjects), were used (Table 3) (*Fig S1*). Details of the stimuli are given in Supplementary Table S1, S2, S3. For sEEG data corresponding to music and movie stimuli, data with >5 inter-ictal spikes were removed from further analysis. The start and end timings of inter-ictal discharges for data with <5 inter-ictal spikes were noted and this segment rejected, with data interpolated using cubic spline interpolation.

#### Electrode localization (Fig 1A)

Intracranial electrodes were localized on a 3-d MNI brain template using a fusion of pre-operative magnetic resonance (MR) and post-operative computed tomography (CT) scans. We started by co-registering the MR T1 image in subject space to standard MNI space (1mm) using the Advanced Normalisation Tools (ANTs; http://stnava.github.io/ANTs/). This process involves fitting, rigid registration, affine registration, and warping using a diffeomorphic field map (SyN algorithm). Next, we used the affine transformation matrix from this co-registration to register the CT image to standard MNI space. The co-registered and normalized CT and MR images were then visually inspected for the anatomical location of the electrode channels (*Fig S1*). The localization of the electrodes in MNI space was corroborated with the implantation maps generated by the epileptologist to clinically localize seizures. The electrodes channels were also assessed on the Harvard-Oxford anatomical atlas (Desikan et al., 2006) to confirm their location in the anatomical regions of interest. The electrode channels from TP and amygdala were then mapped on to the MNI space (*Fig 2 A*)

**Fig 1.**
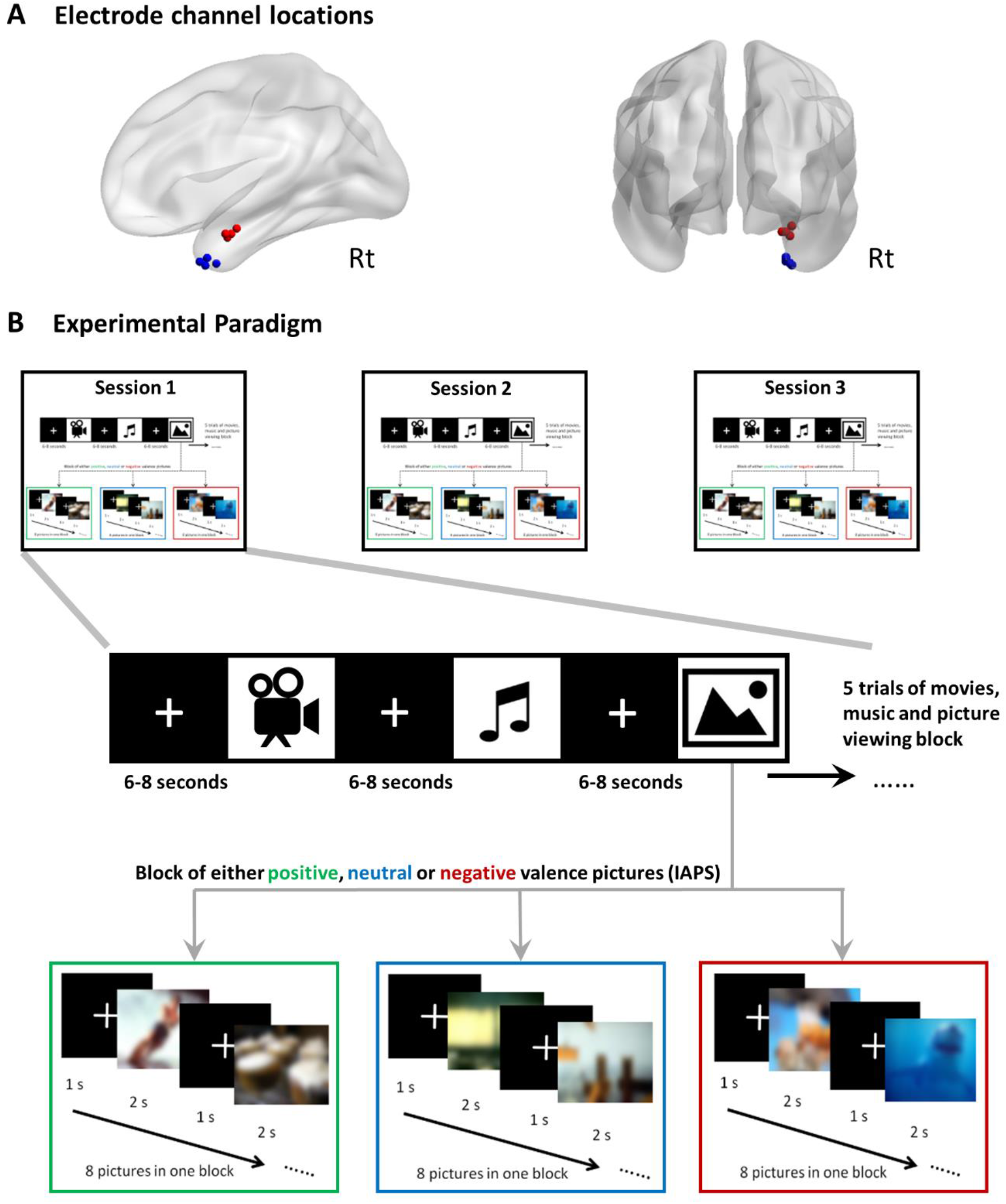
Electrode channel localization and experimental paradigm. **A)** Electrode localization of four subjects (used for data analysis) rendered onto a three-dimensional MNI space. Blue dots indicate electrodes in the temporal pole; red dots denote electrodes in the amygdala. **B)** The experimental paradigm consisted of trials of movie clips (15), music clips (15) and pictures (120) presented in three counter balanced sessions and a break of 5-10 minutes between each session. Each session had five movie clips, five music clips and five blocks of pictures.

**Fig 2.**
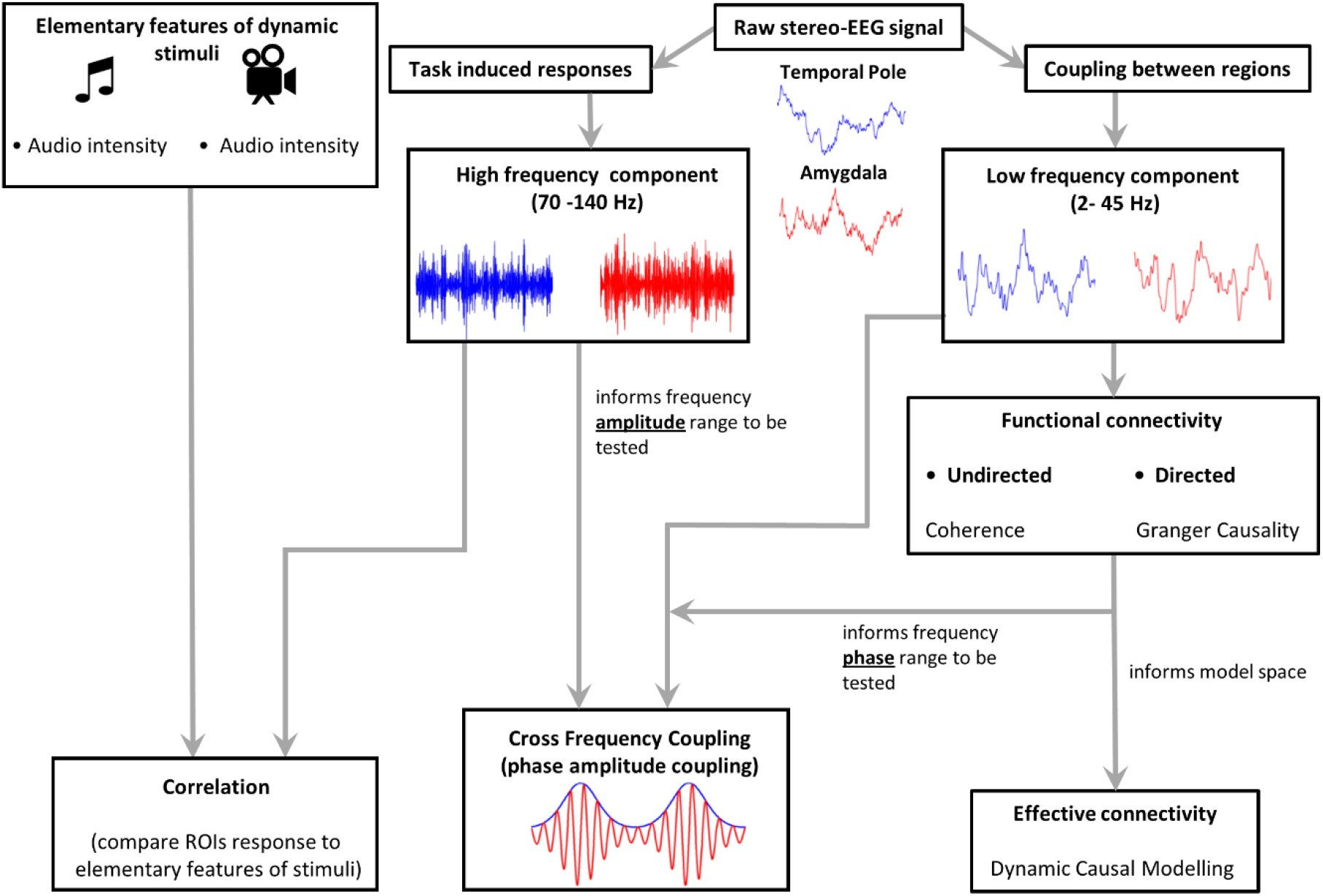
Schema for analytical approach. Hierarchical analysis steps used to uncover the response profile and interaction between the TP and the amygdala activity underlying emotional stimuli.

Based on the above criteria, analyses were performed on one electrode channel pair located in right mesial temporal pole and right amygdala for each subject. Data from P13 was excluded from analysis as frequent inter-ictal activity was present in the data trials. Data from P15 was also excluded because the localisation of the electrode channels were considerably distant (TP: 24, −1, −44, amygdala: 20, −12, −28) from the other participants’ localised channels in the TP and the amygdala (*Table 2*) as lying outside of the target regions according to Harvard-Oxford anatomical atlas.

**Table 2.** Channel location in MNI coordinates.

| Channel | MNI coordinates |  |  |
| --- | --- | --- | --- |
|  | x | y | z |
| TP P2 | 26 | 12 | -38 |
| TP P4 | 28 | 3 | -39 |
| TP P8 | 29 | 9 | -40 |
| TP P10 | 26 | 8 | -36 |
| A P2 | 28 | -03 | -25 |
| A P4 | 29 | -08 | -20 |
| A P8 | 26 | -05 | -23 |
| A P10 | 24 | -02 | -23 |

**Note:** TP – temporal pole, A – amygdala. P2, P4, P8, P10 are subject id.

**Table 3.** Movie clips annotations.

| Clip ID | Mean start time<br>(sec) | SEM<br>(sec) |
| --- | --- | --- |
| **positive valence** |  |  |
| Clip 01 | 69 | 3.43 |
| Clip 08 | 101 | 7.14 |
| Clip 24 | 47 | 2.94 |
| Clip 25 | 46 | 2.80 |
| Clip 31 | 30 | 4.89 |
| Clip 32 | 26 | 2.63 |
| **negative valence** |  |  |
| Clip 02 | 61 | 4.65 |
| Clip 34 | 86 | 7.92 |
| Clip 35 | 36 | 2.11 |
| Clip 40 | 45 | 0.73 |
| Clip 41 | 59 | 4.32 |
| Clip 42 | 44 | 6.68 |

#### Experimental paradigm (Fig 1B)

We employed a paradigm consisting of movie clips, music and static images. There were 3 sessions, each comprising 5 movie clips, 5 music clips and 5 blocks of 8 pictures each. Movie clips were selected according to pilot ratings for valence and arousal by 10 independent healthy participants (*Table S1*, *Fig S2*). Twelve music clips were used from an established dataset rated according to valence and arousal (Eerola and Vuoskoski 2011) (*Fig S2*, *Table S2*). Three music clips were repeated to maintain continuity of the paradigm but only the first presentation of music clips were utilised for our analyses. Static images comprised of emotionally-salient pictures taken from IAPS (International Affective Picture System) database (Lang, Bradley et al. 1997). The sessions were counter-balanced during testing. Participants were presented the stimuli on a laptop with screen size of 14 inches placed approximately 60 cm in front of them. The surgical setting and variable position of the patient in the surgical bed precluded exact control over the distance to the screen. The experimental paradigm was presented using Presentation^®^ software (Version 18.0, Neurobehavioral Systems, Inc., Berkeley, CA, www.neurobs.com).

For both movie clip and music stimuli, a fixation cross appeared at the centre of the screen for 6-8 seconds before the start of the stimulus. The music clips were 14-28 seconds in duration (mean = 18.2 s, *Table S2*) and the movie clips were 50-200 seconds in duration (mean = 86.6 s, *Table S3*). The block of pictures commenced with 6-8 seconds of white fixation cross. Picture stimuli were then presented for 2 seconds each with an inter-stimulus interval with a fixation cross of 1 second. 8 pictures, randomly selected from each respective condition, were presented in each of the blocks (*Fig 3*). There were 5 blocks of pictures for each condition. The standardized set of IAPS pictures has been rated in terms of their ability to induce valence (unpleasant/pleasant) and arousal (calm/excited) changes. Specifically a scale of 1–9 was used for reporting negatively and positively rated emotions. Scores of 1–3 are recognized as negative, 4–6 as neutral, and 7–9 as positive valences. A collection of 32 pictures for positive valence, 37 pictures for negative valence and 31 pictures for neutral valence (NTV) category was first established. Only pictures rated high for arousal were selected for this study (rating 6-9).

**Fig 3.**
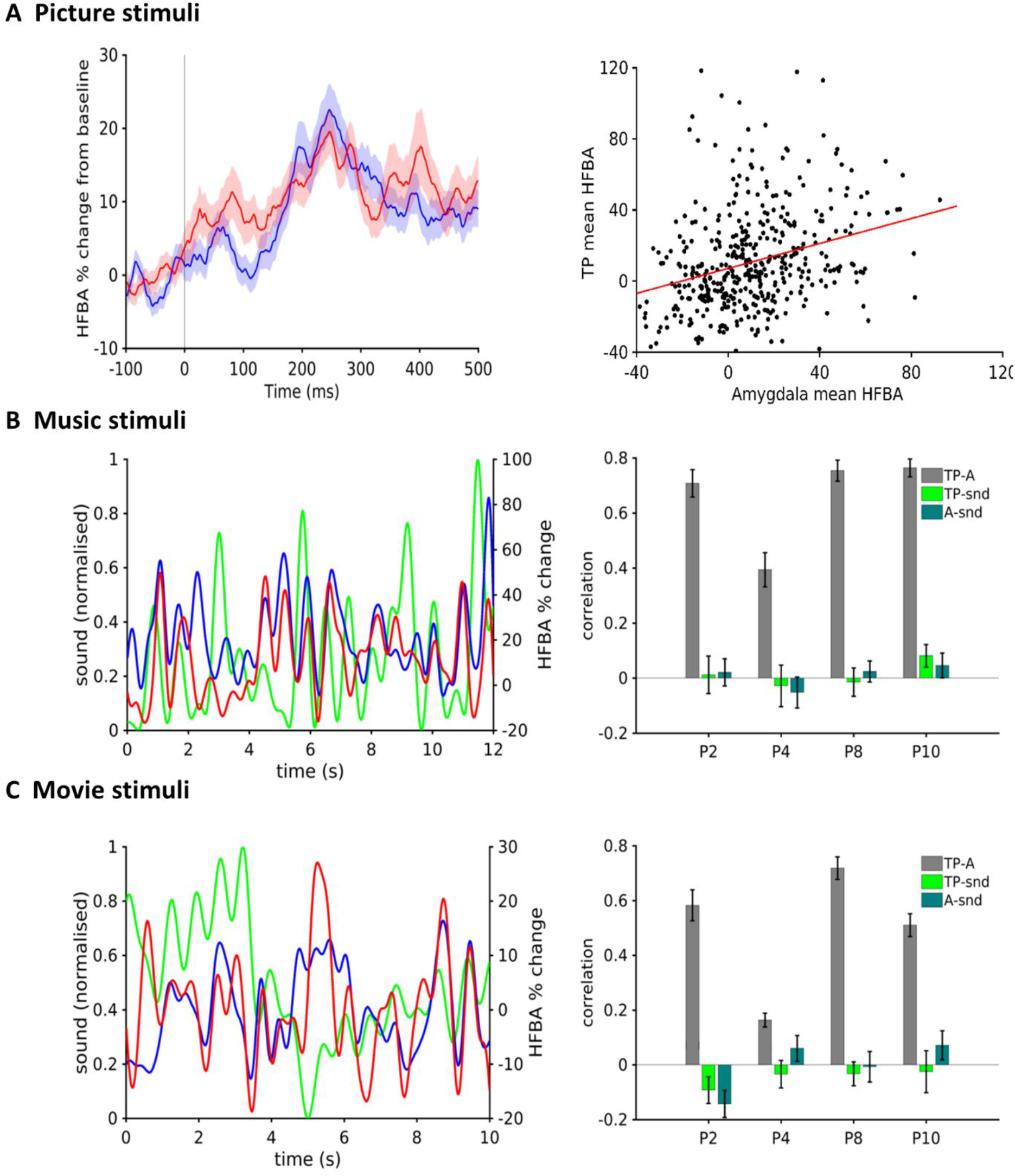
Task induced responses quantified via high frequency activity. **A** Picture stimuli-High frequency activity quantified with high frequency broadband amplitude (HFBA) averaged over all patients and all conditions of pictures for temporal pole (blue) and amygdala (red) (*left*). Correlation analysis for mean HFBA activity for temporal pole and amygdala on a trial-by-trial basis (r = 0.27, *p*<0.0001) (*right*). **B** Music stimuli - sound intensity of one representative music stimuli (green) and corresponding HFBA of the TP (blue) and the amygdala (red) (*left*). Mean correlation between HFBA between TP-amygdala (grey), HFBA TP-sound intensity (light green) and HFBA amygdala-sound intensity (dark green) shown in bar plots on the left (*right*). **C** Movie stimuli-sound intensity of one representative annotated 10 seconds of movie stimuli (green) and corresponding HFBA of TP (blue) and amygdala (red) (*left*). Mean correlation between HFBA TP-amygdala (grey), HFBA TP-sound intensity (light green) and HFBA amygdala-sound intensity (dark green) (*right*). Error bars indicate SEM. Note the high correlation between HFBA of the TP and the amygdala for music and movie clips. P2, P4, P8, P10 are subject IDs.

#### Extraction of elementary acoustic features of naturalistic stimuli

The dynamic sound intensity of music and movie clips was extracted using a Fourier-based spectral density function. Sound intensity was calculated as the average power of the audio derived from the music and movie stimuli. This signal was then smoothed by applying a low pass filter at 0.5 Hz and then downsampled to 10 Hz (Potes, Gunduz et al. 2012).

#### High Frequency Activity (HFA)

We quantified HFA by computing high frequency broadband amplitude (HFBA) as previously reported (Foster, Rangarajan et al. 2015). Briefly, the down-sampled (500 Hz), line removed and re-referenced time series for each channel was filtered between 70 and 140 Hz using sequential band-pass windows of 10 Hz (i.e., 70–80, 80–90, 90–100 … 130-140), via a two-way zero-phase lag FIR filter (EEGLAB). The amplitude (envelope) of each narrow band signal was then calculated from a Hilbert transform. For picture stimuli data, each narrow band time series was expressed as a percentage change from baseline of 100 ms and averaged. For naturalistic stimuli, each narrow band time series was normalised to its mean and expressed as percentage change from the mean.

#### Time varying power spectral density

We computed time varying power spectral density (PSD) of annotated durations of stimuli. We employed a moving window of 1500 ms with an overlap of 1400 ms. The PSD yields information regarding changes in power across different frequency bands also allows testing for ergodicity (i.e. a signal’s time average converges). This validation is important to satisfy the assumptions of inferring effective connectivity analysis via DCM-CSD (*Fig S3*).

#### Coherence

We used the magnitude squared coherence (MSC) as a measure to quantify the functional connectivity between these two regions. Coherence is a measure of the linear relationship of two time-series at a particular frequency and is sensitive to common phase and amplitude dynamics of two signals. MSC is bounded between 0 (no similarity) and 1 (maximum similarity). Magnitude squared coherence was computed as,

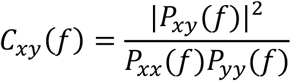

Where *C*_*xy*_ is the magnitude squared coherence between signals *x* and *y*, *P*_*xx*_(*f*) and *P*_*yy*_(*f*) are the power spectral densities of *x* and *y* respectively, and *P*_*xy*_(*f*) is their cross power spectral density.

Coherence was computed using a window of 1000 ms with 500 ms overlap. Coherence values were computed for each epoch and averaged for all stimuli and conditions over the 4 subjects.

#### Spectral Granger causality (spectral GC)

Granger Causality (GC) assesses conditional (asymmetric) dependencies between time series by investigating whether one time series can correctly forecast another (Granger 1969, Bressler and Seth 2011). We used spectral GC, an extension of traditional (time domain) GC in the frequency domain, to characterize the frequency content of directional dependences between the TP and the amygdala. Specifically, spectral GC uses Fourier methods to examine “granger-causality” in the spectral domain (Geweke 1982, Kamiński, Ding et al. 2001) and measures the fraction of the total power at a particular frequency contributed by each signal (self-to-self, self-to-other and vice-versa).

We employed the MVGC toolbox (Barnett and Seth 2014) to compute the spectral GC. The model order *m* was determined according to the Akaike information criterion, which is a trade-off between spectral resolution and complexity. Model orders were estimated for each patient and varied from 7 to 10. The spectral GC curves were then averaged across all the patients. Spectral Granger causality estimates were considered significant if they exceeded the 99.9% confidence interval established by permutation testing (1000 randomisations). The permutation statistics were computed by randomly shuffling the time series in blocks, which take the computed autocovariance lags into account (Barnett and Seth 2014). Duration of each trial was similar as for that of coherence analysis. The spectral GC obtained for each segment of naturalistic stimuli were then averaged for each stimulus and further grand averaged across subjects.

#### Dynamic Causal Modelling (DCM)

DCM provides a computational framework to infer effective connectivity between brain regions from neurophysiological data (Friston, Harrison et al. 2003). DCM employs hypothesis driven model specification and a model inversion procedure based upon variational Bayes. Importantly, DCM allows the effect of experimental conditions to be modelled as modulators of the connectivity between the regions. We employed two types of DCM for event related stimuli and DCM for cross spectral density for naturalistic stimuli respectively. Model space varied slightly in two types of DCM employed for picture stimuli and naturalistic stimuli (see below). We constructed our model space based on all the possible connections between the TP and the amygdala, and all possible manners in which the valence of the stimuli modulated each of these connections. Drawing on the preceding analyses with coherence and sGC, we focussed on the low frequency range (2-12 Hz). For group model comparison, Bayesian model selection (BMS) with random effects was employed to select the winning model at the group level. This approach uses a variational method that to balances the posterior likelihood of a model with its complexity in order to identify the model with the highest exceedance probability (the “winning model”).

#### DCM for Phase coupling (PC) for picture stimuli

We employed DCM-PC, which is based on a framework for coupled oscillators in which the change of phase of one oscillator is influenced by the phase differences between itself and other oscillators (Penny, Litvak et al. 2009). We constructed a model space which encompassed all possible connections between the TP and the amygdala, and all possible manners in which the valence of the stimuli modulated these connections (*Fig 5A*). For picture stimuli, valence modulation was entered as binary values; +1 for positive, 0 for neutral and −1 for negative pictures. Based on possible connections in the model space, DCM-PC can suggest a master-slave relationship in which the master region enslaves the phase of the passive region in a unidirectional model, or a bidirectional model in which a mutual entrainment occurs (Penny, Litvak et al. 2009).

#### DCM-cross spectral density (CSD) for naturalistic stimuli

DCM-CSD was developed and implemented in 2012 (Friston, Bastos et al. 2012), is a generalisation of DCM for steady state responses that has been well-validated across many studies (Moran, Kiebel et al. 2007, Moran, Stephan et al. 2009, Moran, Jung et al. 2011, Moran, Mallet et al. 2011, Moran, Symmonds et al. 2011). Cross spectral density analysis allows one to determine the statistical dependencies between two time series as a function of frequency. To do this DCM-CSD optimises (i.e. fits parameters) to the Fourier transform of the cross-correlation function. The model itself is based on plausible synaptic dynamics that embody excitatory and inhibitory cells and post-synaptic potentials. We used the so-called canonical microcircuit model that incorporates local (within-region) as well as between region connectivity drives (Bastos, Usrey et al. 2012). Crucially our comparison focused on the region-to-region (feedforward and feedback) connections and their modulation by emotional valence. The model space for DCM-CSD encompassed all possible (forward and backward) connections between the TP and the amygdala, and all possible manners in which the valence of the stimuli modulated these connections. The models for feedforward connections are identical to those used in DCM-PC (*Fig 5A*). Feedback connections were also included in the model space but are not illustrated for simplicity. DCM-CSD analysis was undertaken separately for music and movie clip stimuli. For naturalistic stimuli valence modulation was entered as binary values; +1 for positive and −1 for negative stimuli.

#### Cross-Frequency Coupling (CFC) via Phase-Amplitude Coupling (PAC)

To compute PAC, we followed the parametric approach validated previously (van Wijk, Jha et al. 2015). Low- and high-frequency components of the sEEG signals were obtained by bandpass-filtering around centre frequencies between 2 and 12 Hz with 0.5 Hz steps (filter bandwidth ±1 Hz), and between 70 and 140 Hz in steps of 2 Hz (filter bandwidth ±35 Hz). Subsequently we extracted the instantaneous phase for each low-frequency component of TP and the amplitude of each high-frequency component of the amygdala. This was computed by bandpass filtering the signal in the desired frequency range and then using the Hilbert transform to derive the instantaneous phase and amplitude. PAC was then estimated for each low-/high-frequency pair of frequency combinations using a general linear model (GLM). This approach is based on parametric statistical analysis and as such is both computationally efficient and highly sensitive (van Wijk, Jha et al. 2015). The duration of each trial was the same as that used in coherence and GC analysis.

## Results

After exclusion of participants due to ictal discharges in the data and inconsistent electrode placement location, data from 4 participants (see Methods) were subjected to a nested series of analytical approaches (*Fig 2*) to quantify the local responses in each structure as well as statistical and dynamic interdependences between them.

### Local changes in high frequency activity (HFA)

HFA (70-140 Hz) is an index of local cortical processing typically reported in sensory and motor cortices and also observed in higher order regions (Jerbi, Ossandon et al. 2009, Miller, Zanos et al. 2009, Crone, Korzeniewska et al. 2011, Lachaux, Axmacher et al. 2012). We quantified HFA by computing high frequency broadband amplitude (HFBA) as previously reported (Foster, Rangarajan et al. 2015). The TP and the amygdala exhibited similar HFA profiles to the picture stimuli (*Fig 3A left*). Mean peak responses (100-400 ms) of both of these regions were significantly correlated (r = 0.27, *p*<0.0001) (*Fig 3A right*).

The presentation of naturalistic stimuli of music and movies also induced highly correlated fluctuations in HFA between the TP and the amygdala. To clarify whether this reflected common drive by the elementary audio features of the music and movie stimuli, as previously reported (Potes, Gunduz et al. 2012), we computed the correlation between HFA and dynamic audio intensity of the stimuli. HFA from both these regions was minimally correlated to the sound intensity of the stimuli (*Fig 3B, C*) suggesting that the common HFA in the TP and the amygdala likely reflect more complex process induced, but, not driven, by the audio-visual stimuli.

### Functional connectivity between the TP and amygdala

We next used coherence, an undirected measure of functional connectivity to further characterize these statistical dependences. For all 3 stimuli types, responses between the TP and amygdala were coherent across a broad peak centred at low frequencies (~3-5 Hz), extending up to 10-12 Hz (*Fig 4A*). To investigate if this pattern was observed exclusively in response to our experimental stimuli, we computed coherence for the baseline condition of fixation cross. Mean coherence estimates for baseline and task condition did not significantly differ (*Fig 4A*) suggesting an intrinsic pattern of statistical dependences across low to moderate frequencies that are not modified by the presentation of emotional stimuli, despite clear task responses in HFA.

**Fig 4.**
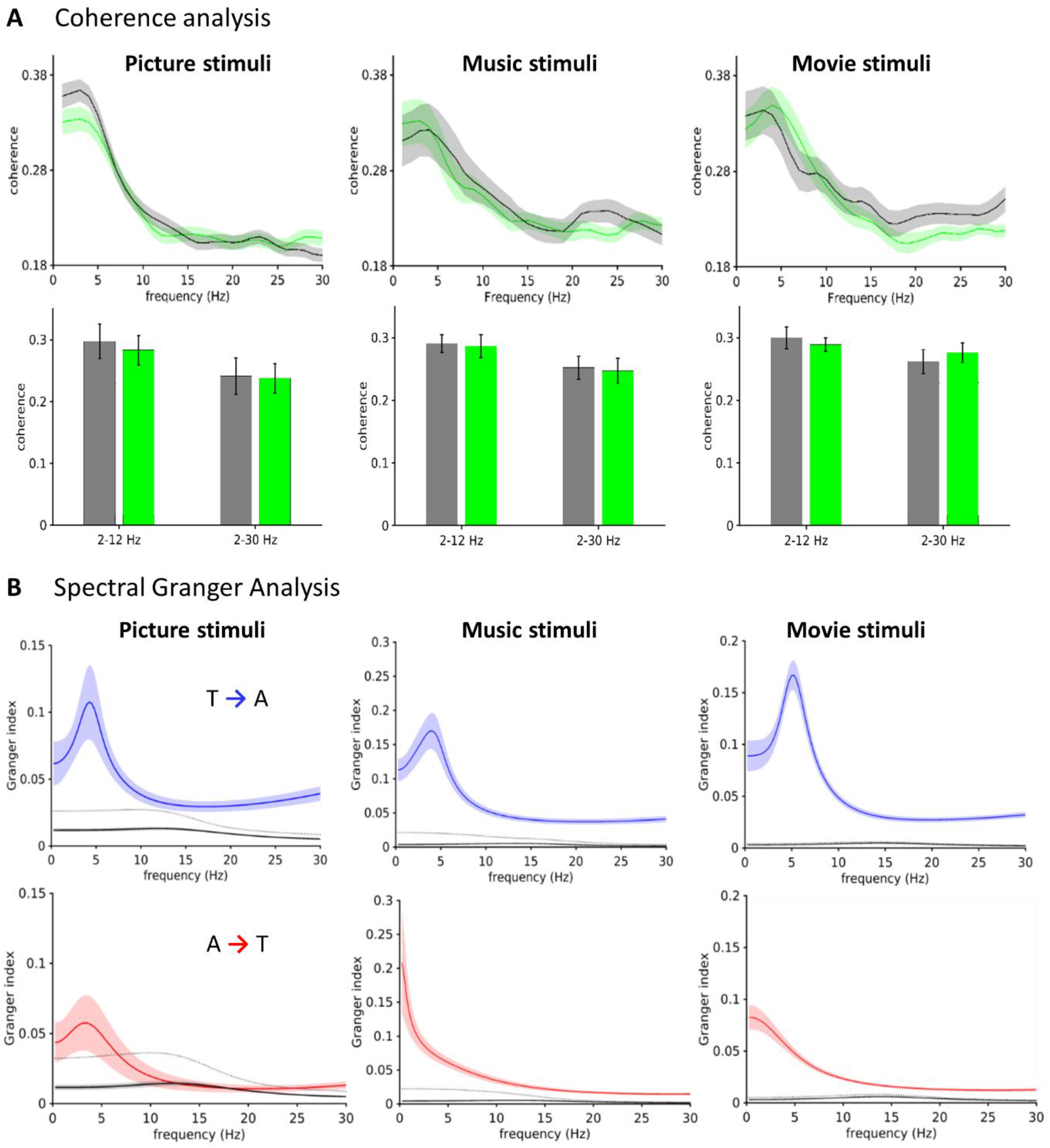
Functional connectivity results. **A** Upper panel-Grand average coherence analysis between temporal pole and amygdala for all conditions—pictures (left), music stimuli (middle) and movie clip stimuli (right). Task (green) and baseline condition (grey) (*left)*. Lower panel-Mean coherence values for two frequency ranges showing non-significant difference tested using a Wilcoxon-ranked sum test (p>0.05) for all tasks—pictures (left), music stimuli (middle) and movie clip stimuli (bottom) (*right*). **B** Upper panel-Grand average spectral Granger Causality showing predominant directed connectivity from the temporal pole (T) to the amygdala (A) for all conditions—pictures stimuli (*left*), music stimuli (*middle*) and movie clip stimuli (*right*). Lower panel-Grand average spectral Granger Causality showing directed connectivity from the amygdala to the TP. These results show peaks in frequency range on 2-12 Hz. Error bars and shading indicate SEM. Black dotted lines denote 99.9% confidence interval obtained via permutation testing with 1000 randomisations and the corresponding grey shading indicates SEM.

To evaluate possible asymmetries in the functional connectivity between the TP and the amygdala, we used spectral Granger causality (GC) analysis - a broadly used measure to infer directed influence between the components of a system (Geweke 1982, Kamiński, Ding et al. 2001). Employing spectral GC to our data revealed a highly asymmetric spectral GC effect from the TP to the amygdala, predominantly in the frequency range of 2-12 Hz, peaking at approximately 3-6 Hz (*Fig 4B*), occurring consistently across all stimuli types. There was a weaker effect in the opposing direction, namely from the amygdala to the TP mostly for picture stimuli and a significant yet attenuated effect in the extremely low frequency range of 1-2 Hz for naturalistic stimuli (*Fig 4B*). Statistical significance was tested at the 99.9 percent confidence intervals obtained using permutation testing.

Spectral effects observed in the TP-amygdala GC analyses were consistent with the frequency range of the coherence. These results thus highlight low frequency contributions to the dependences between these two regions.

We next used DCM-CSD, a generalisation of DCM for steady state responses which allows one to infer frequency-specific directed influences (Friston, Bastos et al. 2012). Application of this DCM rests upon assumptions of ergodicity for longer time segment data. Analyses of time varying PSD in our data showed clear temporal mixing of the power in time (i.e. lack of any clear drift) (*Fig S2*) satisfying the ergodicity assumptions required for DCM.

We found that the DCM-CSD model with the highest exceedance probability for both music and movie stimuli was such that affect modulated a unidirectional connection from the TP to the amygdala (*Fig 5B, C*).

**Fig 5.**
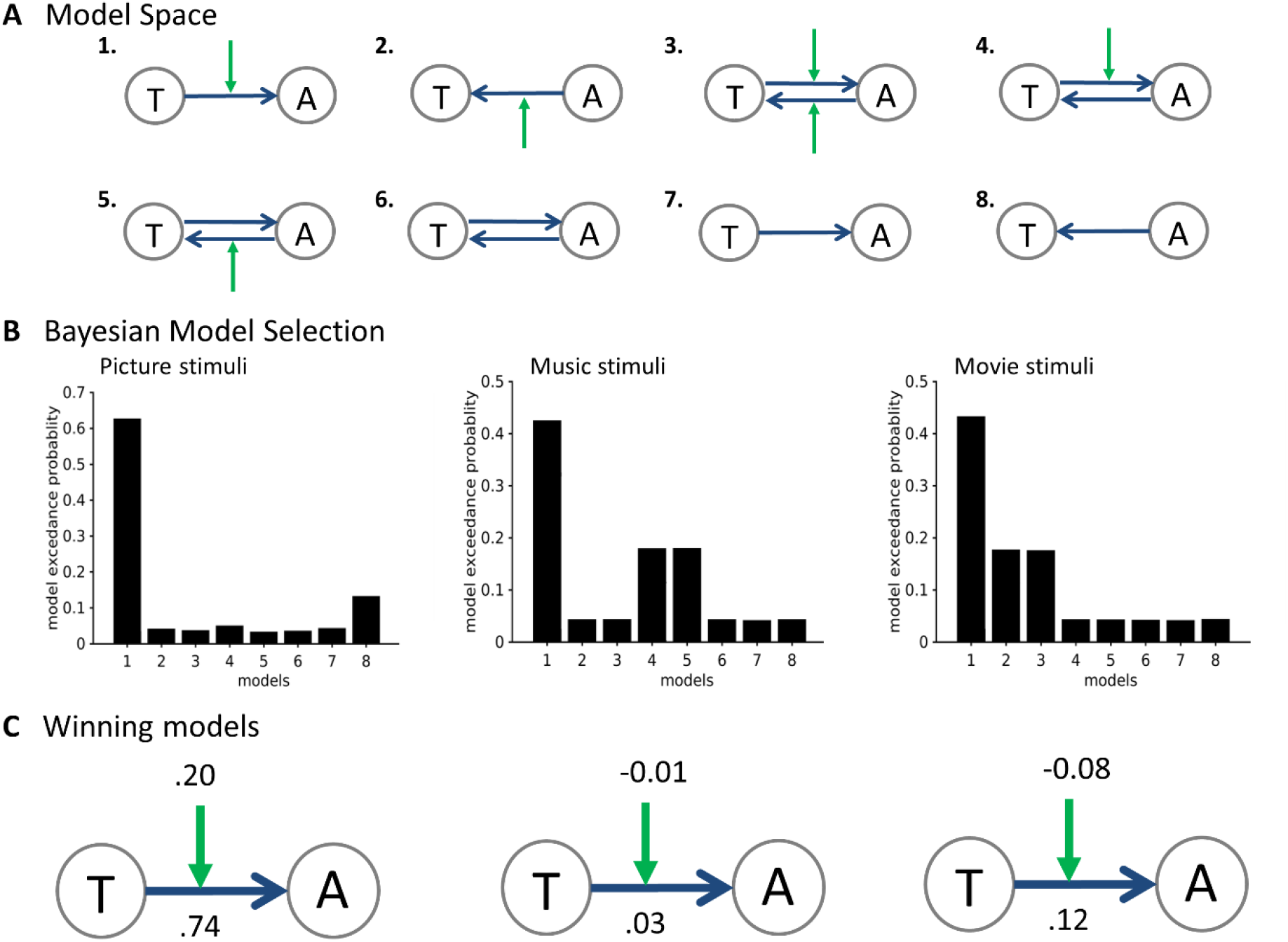
Dynamic causal modelling. **A** The model space comprised eight models, with all possible connections and task modulation effects influencing the connectivity between temporal pole (T) and amygdala (A). Only forward connections (for DCM cross spectral density) are shown for simplicity. Green arrow indicates task modulation (positive, neutral and negative valence pictures and positive and negative valence dynamic stimuli for music and movie stimuli). **B** Bayesian model selection identified model 1 as possessing the highest exceedance probability for DCM employed for all three types of stimuli— picture stimuli (*left*), music stimuli (*middle*) and movie stimuli (*right*). **C** Connectivity and modulatory effects together with the parameter estimates are presented for the winning model in three types of stimuli—picture stimuli (*left*), (ii) music stimuli (*middle*) and movie stimuli (*right*).

### Cross frequency analyses

These preceding results show the presence of local responses in these regions via coherent changes in HFA, together with functional and effective connectivity in low frequencies, specifically in the direction from the TP to amygdala. We next asked whether there was a statistical dependence between low frequency activity in the TP and high frequency activity in the amygdala using cross-frequency coupling (CFC) analysis (Jensen and Colgin 2007, Canolty and Knight 2010). We employed phase-amplitude coupling (PAC) which is a type of CFC that captures a relationship between the low-frequency phase of one neurophysiological signal and the high-frequency amplitude (or power) of the same or another signal (Fries 2005, Fries 2015). PAC analysis revealed that in each participant, the phases of alpha/theta oscillations in TP were significantly correlated with the amplitude of high gamma activity of amygdala (*Fig 6*). The precise frequency-dependent nature of coupling between low and high frequencies showed considerable inter-subject variability.

**Fig 6.**
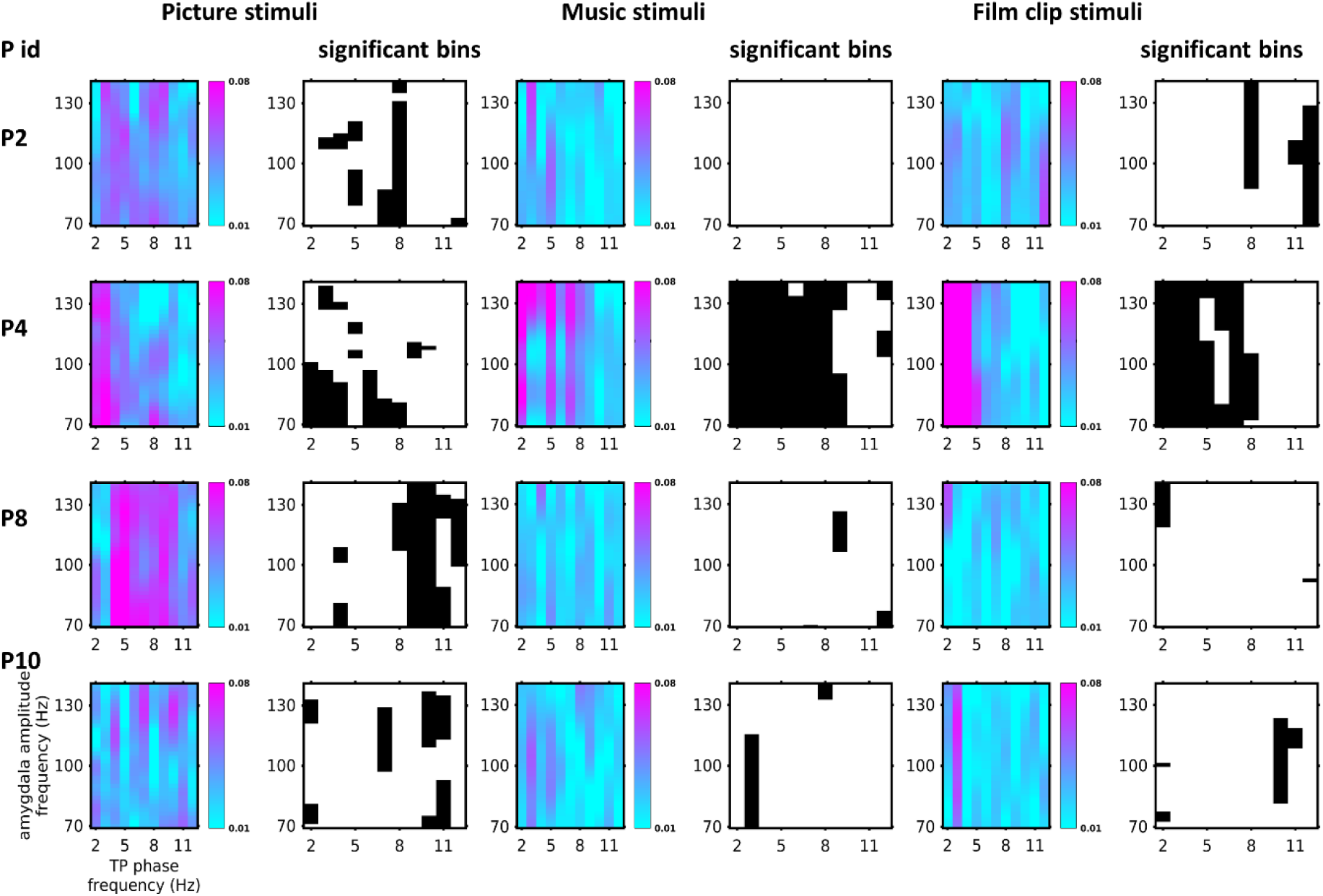
Cross frequency coupling analysis. Phase amplitude coupling (PAC) co-modulogram plots from all 4 participants for all types of stimuli. Each participant for each condition has two plots show—one for showing PAC in arbitrary units and a black and white plot showing significant coupling. The first two columns show PAC for picture stimuli, middle two columns for music stimuli and last two columns for movie stimuli. The high gamma amplitude in the amygdala was phase-locked to the phase of temporal pole low-frequency (theta and alpha) rhythms for all subjects except for music stimuli data for P2. Colour bar scale is in arbitrary units. Binary scale plots show the significant bins in black (*p*<0.05). Some inter-subject and inter-task variability is observed. P id are subject ids.

## Discussion

Using intracranial human EEG, we demonstrate that perception of salient emotional stimuli evokes a complex orchestration of responses in the TP and the amygdala for static and dynamic visual and auditory stimuli. Both regions share a consistent and temporally concordant high frequency response to affective stimuli. These high frequency responses do not correlate with the low-level acoustic and visual attributes of the naturalistic stimuli, suggesting a role in higher order facets of natural stimulus perception such as their semantic or narrative content (Hasson 2004). Undirected and directed functional connectivity analyses for all the stimuli suggest that the interactions between these two regions are mediated by low frequency activity of the TP entraining high frequency activity of the amygdala via phase-amplitude coherence. Effective connectivity analyses suggest that the direction of this coupling was from the TP to the amygdala, with the emotional valence of the stimuli modulating this connectivity.

The apex role of the TP in the TP-amygdala dyad occurs within a broader, embedded role for this region. The TP is association cortex, considered an apical point of the auditory and ventral visual stream. The ventral visual stream has been implicated in mapping perceptual information onto meaning (Ungerleider and Haxby 1994). In principle, neural activity (evoked by stimuli) integrates activity in lower-level visual areas with hierarchical cognitive areas like fusiform gyrus, peaking in the TP and yielding increasing levels of processing complexity (Mishkin and Ungerleider 1982, Moran, Mufson et al. 1987, Felleman and Van 1991, Webster, Ungerleider et al. 1991, Suzuki and Amaral 1994, Saleem and Tanaka 1996). An intriguing study recently supported this functional–anatomical dissociation. Multi-variate pattern analysis suggested that posterior areas of this stream process perceptual properties of objects, whereas conceptual properties (e.g., how an object is used) are processed more anteriorly in the ventral temporal lobe (Peelen and Caramazza 2012). Emotional valence is also a complex conceptual property and the TP has been shown to identify and represent affect as well (Blair, Morris et al. 1999, Royet, Zald et al. 2000, Chikazoe, Lee et al. 2014). Emotional deficits are also reported in patients with lesioned TP (Eslinger, Moore et al. 2011) suggestive of valence processing at the TP.

The crucial role of the TP in emotional differentiation does not exclude valence differentiation by the amygdala. A strong body of research supports the amygdala’s role in emotional valence. For example, animal studies suggest the existence of subsets of basolateral amygdala (BLA) neurons encoding positive or negative valence (Gore, Schwartz et al. 2015, Namburi, Al-Hasani et al. 2016, Beyeler, Chang et al. 2018, Tye 2018). However, auditory (dorsal) and visual (ventral) association regions in the TP send projections to the basal lateral and basal medial nuclei of the amygdala (Ghashghaei and Barbas 2002) which could convey valence related information. Another study also suggests the TP directly projecting to amygdala but not vice-versa (Nieuwenhuys, Voogd et al. 2007).

An additional role of the amygdala is influencing physiological changes, which are integral to emotion experience (and thus also encoding valence) (Cannon 1927, Seth and Friston 2016), may perhaps disambiguate it from the TP. The amygdala has considerable anatomical connections to the hypothalamus and brainstem, and produces visceral correlates of emotional arousal such as heart rate changes (LeDoux 2000). Human neuroimaging has also found association between amygdala activation and measures of autonomic arousal such as skin conductance, blood pressure and heart rate (LeDoux 2000, Yang, Simmons et al. 2007). Amygdala stimulation has also consistently evoked autonomic changes (Mangina and Beuzeron-Mangina 1996, Lanteaume, Khalfa et al. 2007, Inman, Bijanki et al. 2018). Thus, overall it seems likely that the TP and amygdala may have complimentary functions contributing to emotional valence generation.

In addition, emotional parsing to the amygdala via the TP does not preclude the role of amygdala in rapid threat related responses. Specifically, the conditional role of the amygdala in our data is consistent with the “high and low road” hypothesis (LeDoux 1998, Freese and Amaral 2005, Freese and Amaral 2006). Accordingly, the high road traverses from lower order regions to higher order cortices (typically visual cortex (V1) to infero-temporal and area TE before the amygdala) and can encompass conscious perception. Our study positions the TP in the “high road” pathway. On the low road, the superior colliculus directly mediates stimulus responses in the amygdala. For threatening stimuli, the processing of stimuli may be expedited, to allow for a rapid preparation of adaptive responses including physiological changes, with amygdala at the apex of such processing (Morris, Öhman et al. 1999, Carr 2015, McFadyen, Mermillod et al. 2017).

In this study, we employed a conceptual hierarchy of complimentary methods and models to study responses and interactions between the TP and the amygdala. Stereo-EEG channels capture local field potentials (LFP) and are thus less vulnerable to the influence of volume conduction than macroscopic scalp EEG signals (Zaveri, Duckrow et al. 2009). We utilised findings from the coherence analysis and spectral GC to restrict the frequency range to be tested for DCM. Such an approach, where GC provides prior knowledge to constrain model space of DCM, has been previously advocated (Seth and Friston 2016). Notably, the GC and DCM analyses yield convergent results regarding the directionality between the TP and the amygdala. However, DCM-CMC derives from a physiological model which comprises four neuronal populations, comprising spiny stellate cells, superficial pyramidal cells, inhibitory interneurons and deep pyramidal cells (Bastos et al., 2012). Forward connections arise from superficial pyramidal cells, while backward connections arise from deep pyramidal cells (Felleman and Van 1991) corresponding to top-down versus bottom-up hierarchy of cognitive processing (Bastos, Vezoli et al. 2015). Interestingly, the forward connections were stronger for movies which require audio-visual conceptual integration, than music which possesses only unimodal auditory information, again supporting the role of TP as a conceptual store. However, our task simply required passive perception of stimuli: An active task may have elicited more symmetric feedback processes to support prediction and learning (Friston 2005).

Our analyses revealed common high frequency responses and a directed low frequency phase-dependent influence. Phase-amplitude CFC has been observed in various cortical and subcortical sites under various tasks (Spaak, Bonnefond et al. 2012, Fries 2015, Staresina, Bergmann et al. 2015). It has been previously shown that cross-frequency phase amplitude coupling is involved in sensory integration, memory process, and behavioural selection (Lisman and Idiart 1995, Schroeder and Lakatos 2009, Voytek, Canolty et al. 2010). The presence of coupling between the low frequency phase of the TP and the high frequency activity in the amygdala, supports the communication-through-coherence hypothesis (CTC) (Fries 2005, Fries 2015). Briefly, high frequency activity (gamma spikes) arriving at excitability peaks (of the low frequency phase) of the receiving neuronal group promote effective communication and narrows the temporal window to that which supports spike-time dependent plasticity. This putatively promotes efficient communication as task-relevant ‘target’ spikes are given computational salience over distracting inputs (Méndez, Pérez et al. 2014, Voytek and Knight 2015).

There are several caveats of the study. First, inherent effects of epilepsy in our study cannot be ruled out. We utilised the data from non-epileptogenic zones (except one patient) but employed strict quality control of data such as exclusion or data interpolation to mitigate the effects of inter-ictal discharges. Second, the study is limited by a small number of patients. We tested 13 patients with this experimental paradigm but only 6 patients had electrodes in the appropriate locations and, after further exclusion criteria for location and data quality, only 4 were suitable for analysis. However, we used multiple stimulus modalities and complimentary analyses, providing convergent results. Third, the amygdala is composed of various sub-nuclei and we could not precisely locate the channels within specific nuclei. However, high frequency activity seen in our data indicates that the sampled neuronal population were responsive to the stimuli. Finally, we used common average referencing scheme as there is currently no gold standard for the referencing scheme to be used in SEEG. Bipolar referencing in the presence of close placement (1.5 mm) of channels acts as an effective low pass spatial filter with impact on spectral content, particularly of coherence estimates (Michelmann, Treder et al. 2018). Furthermore, the spatial spread of LFP is several millimetres (~3 mm) (Kreiman, Hung et al. 2006, Kajikawa and Schroeder 2011) causing suppression of neural activity from bipolar referenced electrodes. White matter referencing is also an option, but white matter is not electrically inert and affected by nearby as well as distant grey matter (Mercier, Bickel et al. 2017).

In sum, this work highlights a hierarchically super-ordinate influence of the TP over the amygdala. Crucially, these results occurred with all the types of stimuli, including static and naturalistic stimuli. Future work could build on this, and link physiological responses elicited by emotions to the activity and coupling between the TP and the amygdala as well as probing functional hemispheric asymmetry.

## Acknowledgements

We thank Annett Koenig for her assistance in arrangements for data acquisition and Matthew Aburn for helpful discussions. We also thank Jane Stadler for providing the movie clips.

## Supplementary Material

**Fig S1.**
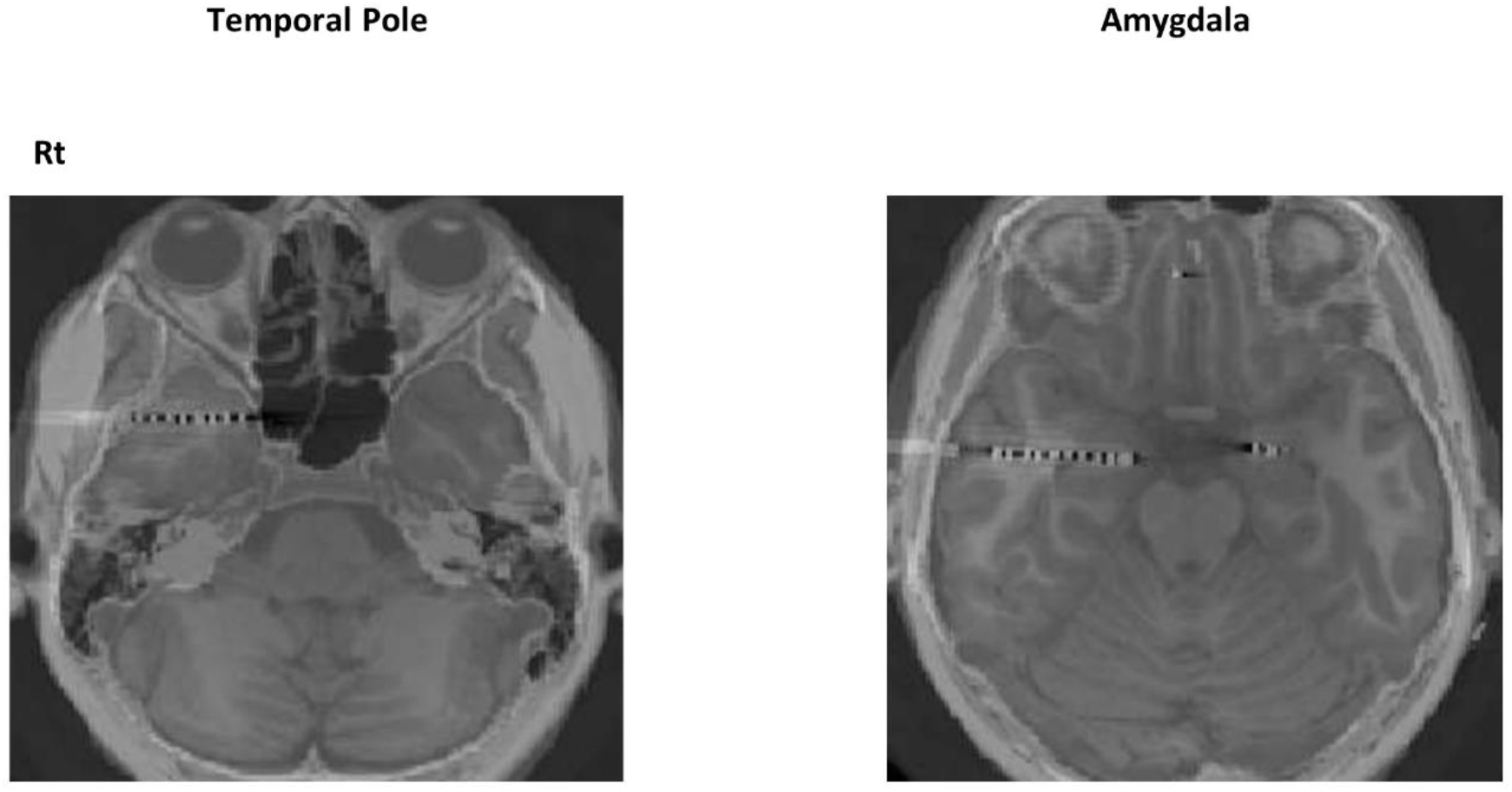
Electrode localization scheme. Electrodes were localized in each participant using co-registered pre-implantation T1-weighted MRI structural scans and post-implantation CT scans **(a)** Example MRI and CT coregistration in MNI space of single subject. The temporal pole electrode (top), the amygdala electrode (bottom). White dots are the electrode channels. Channels in regions of interest were selected based on implantation maps used for clinical evaluation and visual inspection and confirmed with MNI anatomical atlas (Harvard-Oxford). Electrodes in white matter were also not used. Each electrode location was determined by selecting the centre of the electrode and determining which ROI best encompassed the centre of the electrode.Rt-right.

**Fig S2.**
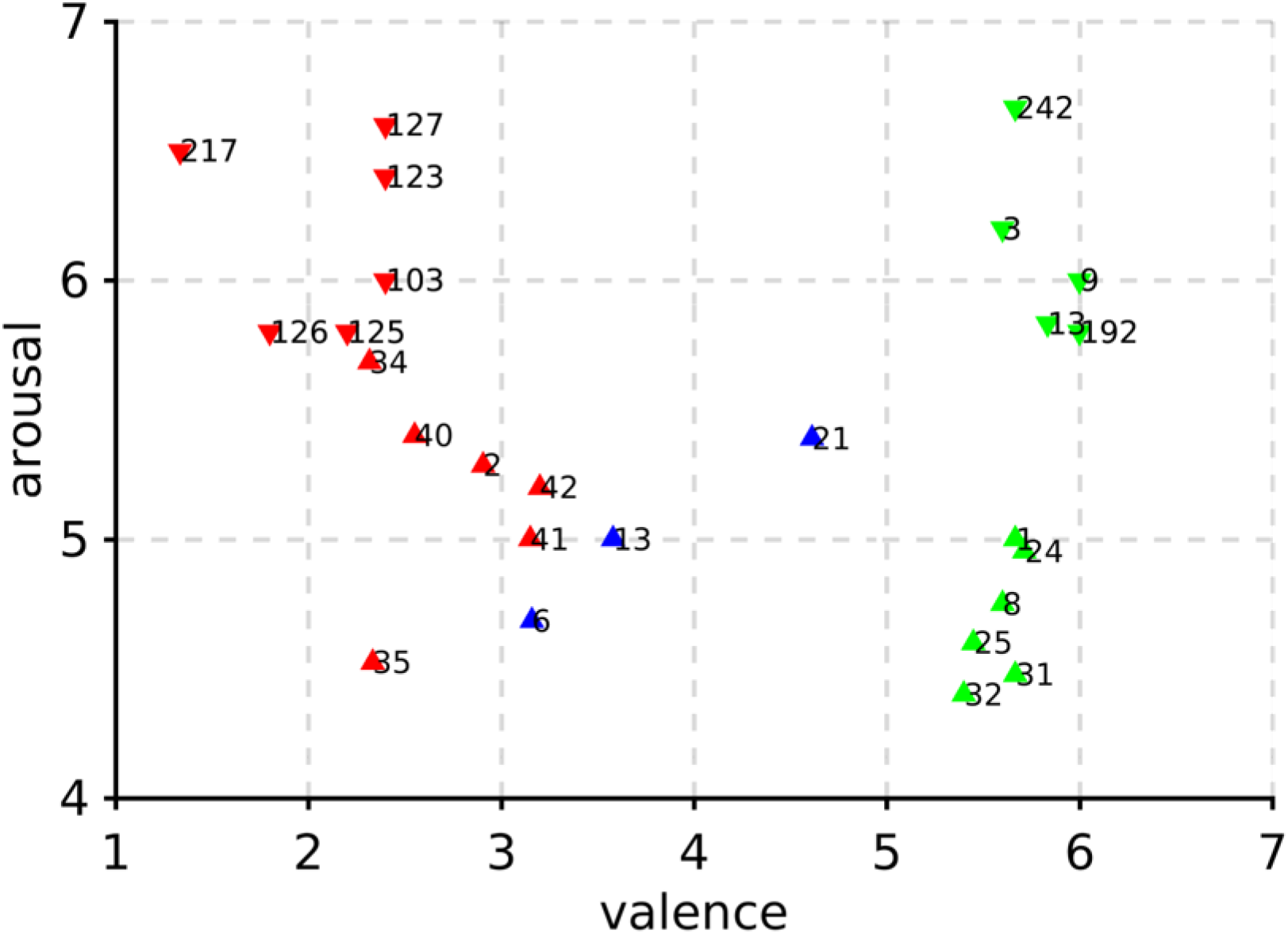
Dimensional scale emotional ratings of naturalistic stimuli. Valence ratings of music stimuli (down pointing triangles) and movie stimuli (up pointing triangles, ids in table S2) categorised them in positive (green), neutral (blue) and negative (red) valence. All stimuli were chosen with high arousal ratings. Numbers beside the data points denote the stimuli id (see *Table S2 and S3*). Neutral valence movie clips were not used for analysis.

**Fig S3.**
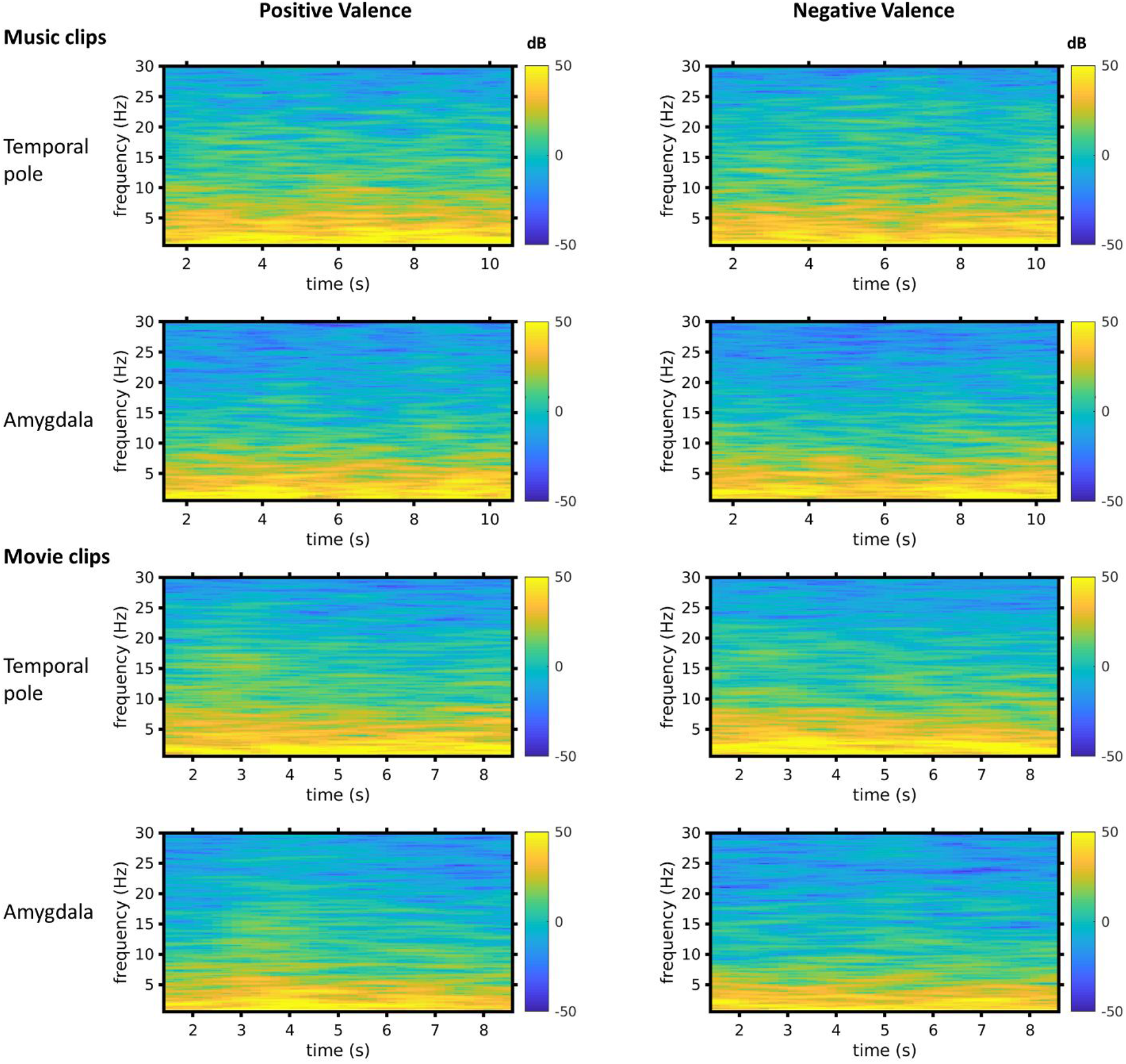
Power spectral density of annotated music and movie stimuli. Average time varying power spectral density for positive and negative stimuli showed most of the power resided in the low frequency range of <15 Hz.

**Table S1:**
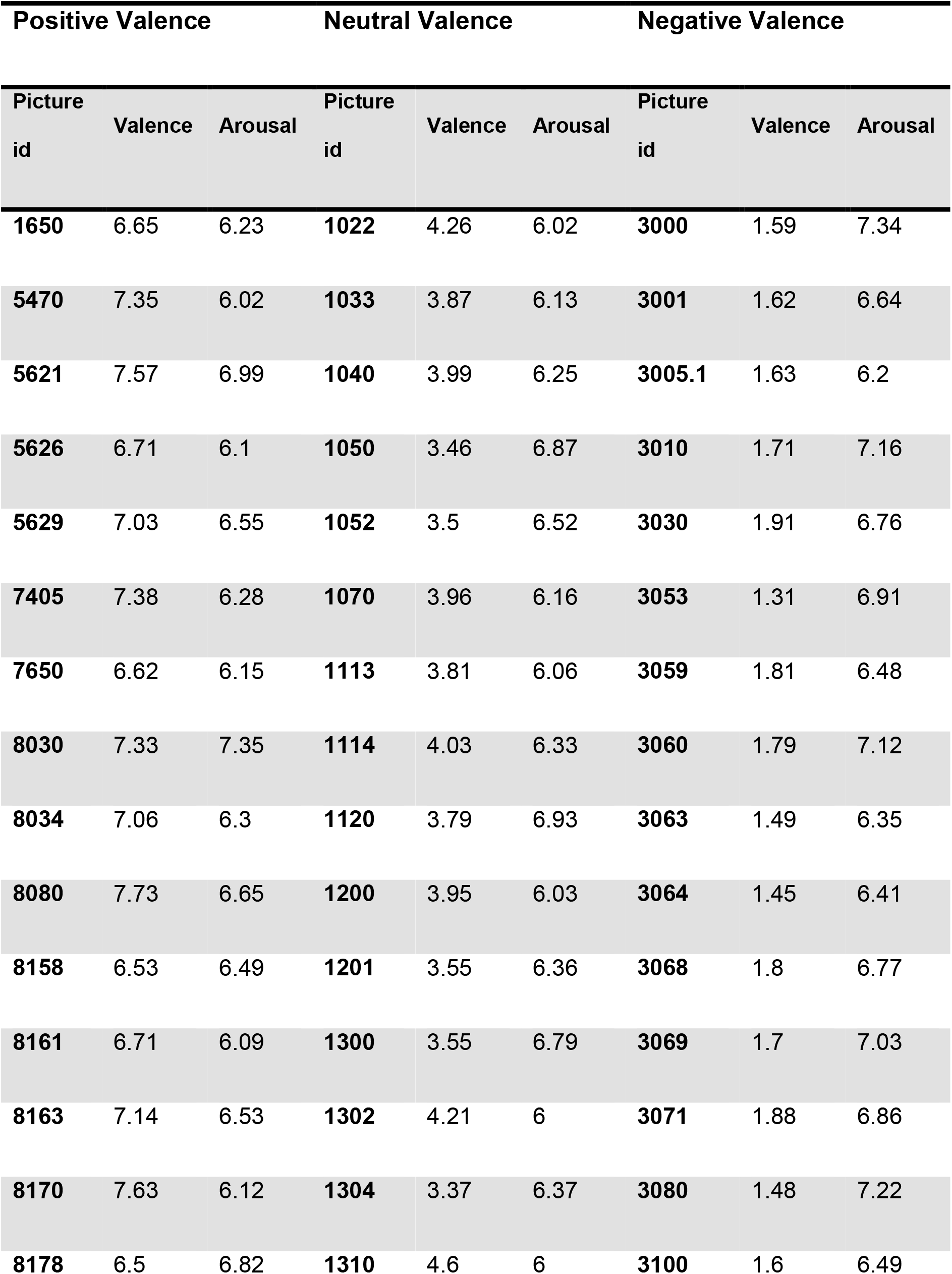
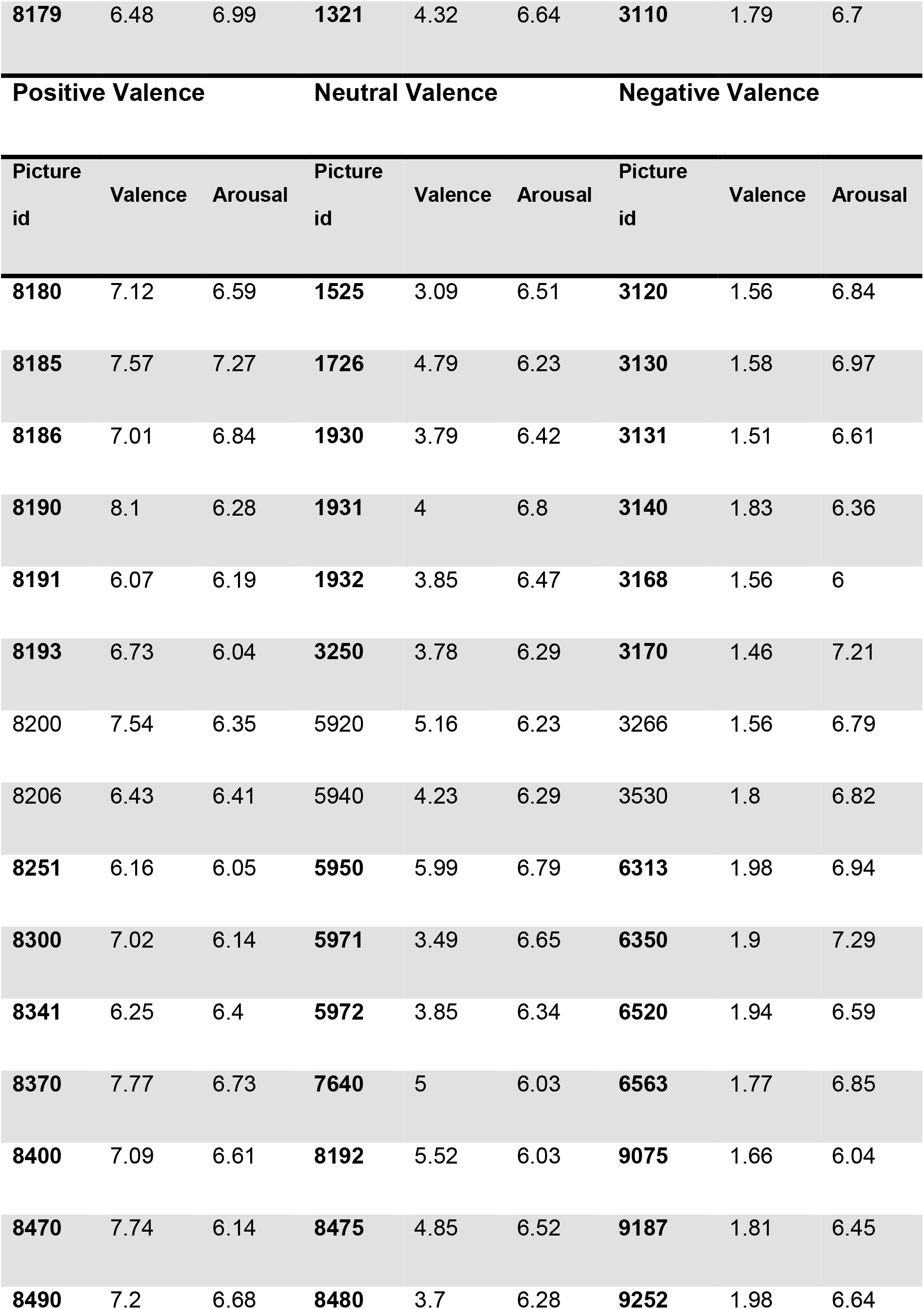
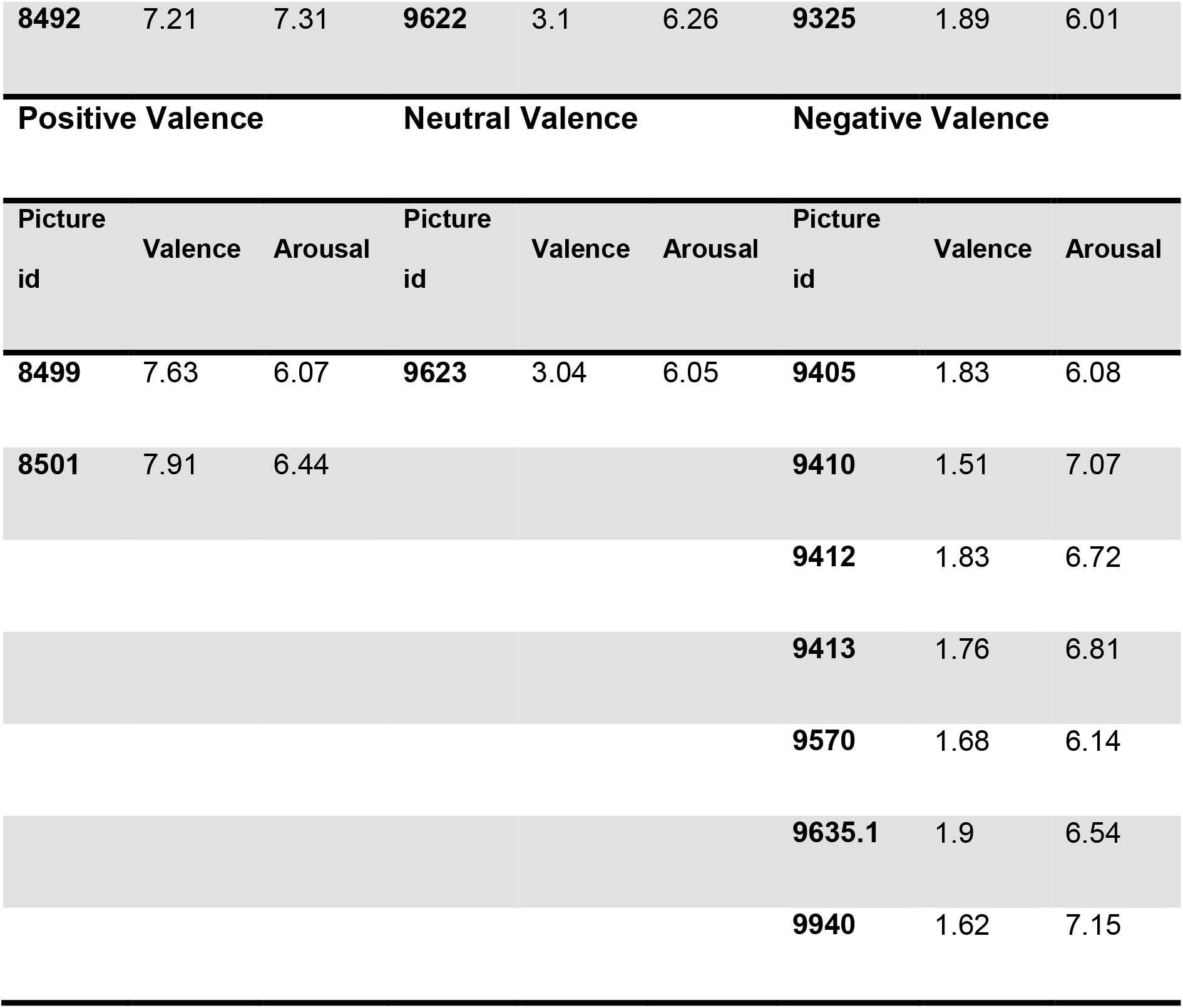
Picture id from IAPS dataset used as stimuli

**Table S2:** Music clips used as stimuli.

| Music clip id | Duration<br>(seconds: milliseconds) |
| --- | --- |
| **Positive Valence** |  |
| 003_HV_HA_Happy | 18.6 |
| 009_HV_HA_Happy | 17.7 |
| 013_HV_HA_Happy | 14.7 |
| 192_HV_HA_Happy | 28.1 |
| 242_HV_HA_Happy | 18.5 |
| **Negative Valence** |  |
| 103_LV_HA_Fear | 15.2 |
| 123_LV_HA_Anger | 21.4 |
| 125_LV_HA_Anger | 16 |
| 126_LV_HA_Anger | 14.7 |
| 127_LV_HA_Anger | 22 |
| 217_LV_HA_Anger | 16.1 |

**Table S2:** Movie clips used as stimuli.

| Movie clip id | Duration<br>(minutes : seconds) |
| --- | --- |
| **Positive Valence** |  |
| Clip 01 10 THINGS I HATE ABOUT YOU song | 1.39 |
| Clip 08 DEAD POETS SOCIETY desk scene | 2.32 |
| Clip 24 Love Actually - All you need is Love Wedding Scene | 1.33 |
| Clip 25 Love Actually - Dancing Prime Minister | 1.03 |
| Clip 31 Mr.Bean s Holiday (Dance Scene) | 0.57 |
| Clip 32 Mr Bean s.Holiday | 1.06 |
| **Neutral Valence** |  |
| Clip 06 ARMAGEDDON sad goodbye | 1.19 |
| Clip 13 GREY THE WOLF CHASE ED | 1.10 |
| Clip 21 KILL BIL VOL1 Vernita Green fight | 1.34 |
| **Negative Valence** |  |
| Clip 02 21 GRAMS hospital grief | 1.43 |
| Clip 34 PARANORMAL ACTIVITY GROWL | 2.05 |
| Clip 35 PINK FLAMINGOS DOG POO | 1.01 |
| Clip 40 REQUIEM FOR A DREAM NEEDLE | 0.56 |
| Clip 41 SHINING THE Here's Johnny | 1.15 |
| Clip 42 Saving Private Ryan short | 1.33 |

